# Larval protein restriction interacts with adult diet to increase fitness in an outbred multiparent population

**DOI:** 10.1101/2024.02.10.579751

**Authors:** Andrew Jones, De’anne Donnell, Elizabeth G. King, Enoch Ng’oma

## Abstract

Developmental conditions including temperature, diet, and parasites can shape adult fitness phenotypes in many species. Studies typically focus on the additive effects of early-life and adult life conditions on life history response in the context of competing models of developmental plasticity (i.e. environmental matching and silver spoon). These models continue to yield mixed results in the same or different species. Here, we characterize interaction effects of larval vs adult diet on lifespan and fecundity in a high diversity outbred population of *Drosophila melanogaster*. We compare fitness proxies of matched vs mismatched early-to-late nutritional conditions differing in protein content. Diet interactions significantly affected both traits, albeit differently. We find no consistent evidence for either model. Rather, several patterns emerged including age and sex effects, survival differences in the post-median life phase, regime-specific timing of peak fecundity, and substantial fecundities in older post-median flies. Surprisingly, lower adult protein delayed egg-laying by about 2 weeks, compared to treatments with higher protein in adult diet. Our results underscore the extent to which adult life history expression depends significantly on developmental conditions. Results further highlight the need for assessing possible adult life phase patterns which might potentially reveal trade-offs between adult life stages than traits.

## Introduction

An organism’s adult phenotype is shaped by a complex interaction of current, developmental, and ancestral environmental conditions including temperature [1–3], food availability [4–6], microbiotas [7,8], or parasites [9]. In particular, nutritional conditions play a critical role in dictating metabolic state dynamics, which impact reproductive, and aging traits in a variety of species from yeast to primates [3,10–12]. The specific schedule of reproduction and longevity an organism takes in current and historical food availability situations is generally thought to arise from optimization of resource allocation to fitness functions. The number of offspring produced can be a function of lifespan, underpinning the quality of somatic tissue maintenance. Much research continues to focus on intergenerational effects of variations in nutritional supply and quality on offspring development and fitness [4,13,14]. Thus, we now appreciate that poor maternal conditions can limit offspring development through poor provisioning and care of eggs and juveniles [15].

The link between developmental diet and adult fitness traits is established in many species [3,16–18], including insects [3,9,16,19], birds [1,5,20,21], fish [18,22], and mammals [17]. For example, Deas et al. [4] found that same generation *D. melanogaster* responded strongly to rich vs poor diet in several morphological traits. However, development time responded strongly to both poor and rich diet, underscoring trait-level variability in diet responses. In another study of *D. melanogaster*, May et al, [23] found that the magnitude of the response at a given stage of life depended on the selection regime experienced in other stages. Similar observations have been made also in cichlid fish with indeterminate growth [18,22]. In the mouth brooding cichlid fish *Simochromis pleurospilus*, Taborsky [18] found that growth in adult fish was dictated by current nutritional availability. Interestingly, like in *D. melanogaster* (Deas et al, 2019), *S. pleurospilus* reproductive rate and progeny size were only influenced by juvenile diet regardless of adult ration. These studies suggest potential trait-level interaction effects of early life, juvenile and adult nutritional conditions.

Early life conditions including nutritional conditions can interact with individual genetic and/or epigenetic states and sex to determine adult fitness. For example, genetic interactions were found to underly larval diet and postponed reproduction in experimentally evolved *D. melanogaster* [24]; male genotype – diet interaction explained some variation in male reproductive traits in isogenic *D. melanogaster* lines [25]; and, larval diet linked with decreased ribosome expression and transcription of certain proteins in lifespan extension [26] in *D. melanogaster*. Other studies demonstrate that early life reactive oxygen species (ROS) mediate stress resistance and lifespan via interaction with ROS-sensitive epigenetic marks in the nematode *Caenorhabditis elegans* [27], and, early experience was linked with structural variation in neuronal genomes in mice [28]. These studies provide support for the role of early diet in adaptive responses to nutritional variability.

The sexes differ in their sensitivity to nutrition during development. For example, if sexual size dimorphism affects survival, the heterogametic sex is more vulnerable to deleterious sex-linked recessive genes if they do not attain optimal size, or high male androgen requirements affects other systems, reviewed in Lindstrom, [5]. Thus, if sexually dimorphic traits are differentially sensitive to nutritional availability, we might predict sex differences in fitness performance. In captive zebra finches, however, a recent study [20] could not find sex-specific environmental sensitivities, although female lifespan was shorter due to senescence, concluding that low food availability shortened lifespan only in individuals developed in harsh conditions. Similarly, in *D. melanogaster*, Klepsatel et al, [3] used a full factorial design to test for differences in flies raised on low-yeast or high-sugar diets and observed reduced reproductive performance regardless of adult conditions. These conclusions are consistent with the silver spoon effect reviewed in Monaghan, [6].

The silver spoon model posits that individuals who develop in favorable conditions will gain fitness benefits throughout their lifespan [29]. Thus, fitness always improves with further improvement in the adult environment; thus, those developed in poor conditions always attain lower fitness compared to those reared in good conditions even with improvement in adult conditions. On the other hand, the environmental matching model predicts that individual fitness will be higher if they experience similar developmental conditions as adults [30]. There are studies in support or disagreement with either hypothesis, although more studies lean towards the silver spoon theory. The SS model has received support in fruit flies [3], birds [20], fish [18]. In the latter, resource allocation to growth and reproduction was shaped by juvenile rather than adult conditions. A meta-analysis of fourteen wild bird and mammal species found a small but significant silver spoon effect of developmental environment effects on survival [21]. Further, May and Zwaan [26] also used a factorial design to vary diet between early life and adulthood, and measured lifespan, fecundity, and gene expression at middle and old age in *D. melanogaster* and found no evidence of mismatches in both lifespan and fecundity responses. What is common to most studies is the focus on understanding the influence of early life conditions on fitness and were performed in relatively fewer genotypes.

In this study, the goal was to characterize the joint effects of larval-to-adult diets (i.e., their interaction) on lifespan and fecundity in a high diversity multiparent population of *D. melanogaster* [31,32] admixed from 835 recombinant inbred lines [33]. Based on the literature reviewed above, we compare fitness outcomes of two matched vs two mismatched early-to-late nutritional conditions differing only in the concentration of a protein source. Specifically, we test the following predictions:

1. Matched diets across larval and adult stages result in higher fitness than if diets are alternated in any sequence (low-high or high-low).
2. Early life diet exerts greater impacts on lifespan and fecundity, such that better (high) developmental diet should lead to higher fitness.

## Methods

### Experimental population

We used a population of *D. melanogaster* derived from the from the *B* sub-population of the *Drosophila* Synthetic Resource Population (DSPR), characterized and described elsewhere [31,32]. Briefly, the p*B* population comprises >800 recombinant inbred lines (RILs) generated from eight inbred founder lines through round robin intercrossing for 50 generations followed by full sibling mating for 25 generations to create inbred lines. For this study, we re-created an outbred population by intercrossing progeny of five females that had mated intra-line, pulled from each of 835 RILs. We let the resulting population randomly mate for five generations. The resulting outbred population was maintained on a standard diet (maintenance diet see below) in six replicates of large cages. We used one of these cages as source of individuals for this study.

### Study design

We implemented a 2 x 2 factorial design (Fig. 1), with two life cycle stages (larvae and adult) and two levels of environmental manipulation: dietary restriction (DR) and control (C). This setup was run in duplicate. To generate experimental individuals, eggs oviposited within a 24-hour duration were collected from the outbred population by slicing out a thin surface layer of media containing 50 - 90 eggs estimated visually and grafting onto a vial surface containing a treatment diet for development on either control or restricted diet (vial type: 25 mm x 95 mm, Polystyrene Reload, cat. No. 32109RL, Genesee Scientific, USA). Emerging flies (12-14 days post-oviposition, po) were released randomly into each of eight cages (dimensions 20.3 cm x 20.3 cm x 20.3 cm) such that flies emerging from the DR and C larval diets are split equally and reciprocally allocated to each of the same diets for adult treatment (Fig. 1). Each cage received a fresh plate of food (100 mm x 15 mm, cat. No. FB0875713, Fisher Scientific, USA) three times a week (Monday, Wednesday, Friday). In addition, each cage received a separate micro-plate (60 mm x 15 mm, cat. No. FB0875713A, Fisher Scientific, USA) containing moist cotton wool as an additional source for drinking.

**Figure 1:**
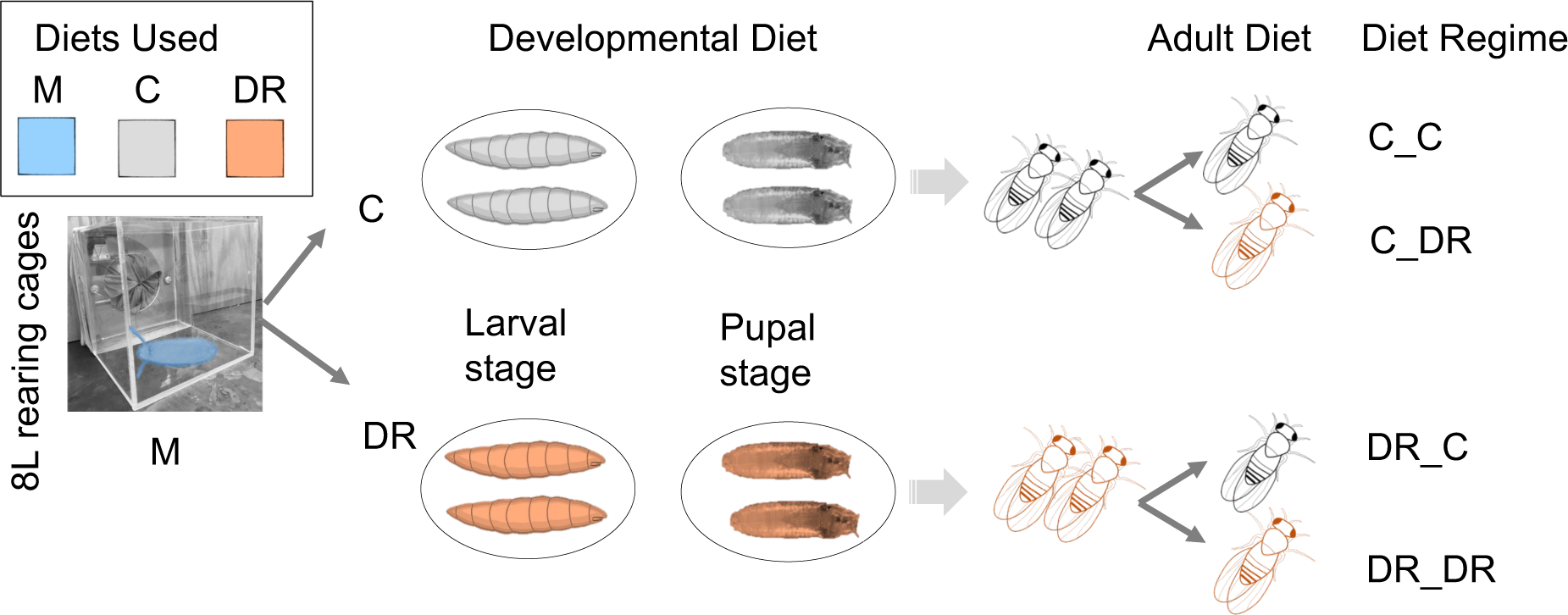
A factorial study design. Eggs collected from an outbred population reared on maintenance diet (M) were seeded on C and DR diets for development. Emerged flies from each treatment were split to C and DR diets as adults in two large cage replicates for each treatment (N_CAGES_ = 8). From each cage we measured fecundity and lifespan.

### Diet description

We maintained RILs used to generate the outbred population (hereafter, OP), and the OP itself on a cornmeal-dextrose-yeast maintenance diet (hereafter, M) containing estimated protein (P, from yeast and cornmeal) to carbohydrate (C, from yeast, dextrose, and cornmeal) proportions of 0.12. We used two mild variations of sucrose-yeast experimental diets with calculated P:C ratios of 0.63 for the DR diet and 0.89 in the C diet. All diets used inactivated SAFPro Relax + YF 73050 yeast brand (Lesaffre Yeast Corp., Milwaukee, USA) which typically provide 45 - 60gprotein and 30 - 38g carbohydrate per 100g yeast. Food values from yeast are calculated using the mid value of the range, whereas those for dextrose, sucrose and cornmeal are FDC values from USDA, (published: 4/1/2019, accessed June 21, 2020; terms: cornmeal, whole-grain, yellow; granulated sugars, sucrose; and powdered sugars, dextrose). All prepared diets were stored at 4 ℃ and used within 14 days of preparation. Each cage received a fresh plate of food every Monday, Wednesday, and Friday, and with each egg quantification event. We reared experimental flies in a temperature-controlled chamber at 23 ℃, 24:0 light-dark cycles, which are typical rearing conditions for the DSPR.

### Phenotype measurement

We measured two life history phenotypes: lifespan, as number of days an individual lived from eclosion until death or censoring, and fecundity, as the number of eggs oviposited in a 3-hour period, three times a week (Monday, Wednesday, and Friday) consistently at the same time of day (12 noon to 3 pm). To measure lifespan, we counted the number of dead flies in each cage by sex, noting censored individuals (escaping or killed accidentally) whenever possible. Food plates bearing eggs for fecundity estimation were stored in a -20 ℃ freezer until enumeration using approach described in [34,35]. Briefly, we filtered eggs from the media onto black fabric discs using a custom vacuum suction pump and took images in a light-controlled chamber with a Canon Rebel TS*i*, (Canon Inc, Japan) camera at exposure time 1/25 s, aperture F5.6, and ISO 100. Eggs on one disc represented collective fecundity of females in a single cage for a period of 3 hours. We used ImageJ software [36,37] to count eggs from each image manually by clicking over eggs and the tally exported to a comma separated value (csv) file. Using the record of death and censoring on each phenotyping date, we calculated average age-specific female fecundity three times a week until survival reached zero. All data, including lifespan records, egg counts, and original egg images were uploaded to and can be accessed from https://zenodo.org/records/10445786.

### Statistical analysis

#### Survival

We explored patterns of survival in each diet regime (L_C__A_C_, L_C__A_DR_, L_DR__A_C_ and L_DR__A_DR_) using Kaplan-Meier survival models and compared survival rates based on log rank tests using the Survival package in R environment [38]. With the aim to understand longitudinal trends in survival, we analyzed data at 10 % and 90% in addition to the conventional 50% survival (i.e. median lifespan). In population studies 10% survival is the mean lifespan of the longest-lived 10% of a given cohort of a species, see [39,40]. Generally, we expect non-stochastic early mortality to stabilize near 95% making 90% a reasonable level to record biologically driven mortality. On the other hand, 10% survival is anticipated to represent maximal longevity at population level.

Our exploratory analysis suggested temporal differences in survival rates associated with both treatment and sex differences. We therefore asked if the risk of adult death differed among treatment groups as a function of larval diet. We therefore constructed a Cox model to compare the baseline hazard function while accounting for time to understand the effect of covariates (larval-adult diet, sex, and cage) on the survival function. In a Cox model, the risk of death is assumed to be proportional if the effects of all covariates do not change over time. Because larval diet and sex violated this assumption, we stratified the model on sex (the variable violating the assumption most), to allow the two strata (male, female) to assume different baseline hazards, and coefficients to be constant across strata [41,42]. This strategy effectively corrected global test statistic (χ^2^ = 4.635, *p* > 0.05) and the test for larval diet only marginally (χ^2^ = 3.509, *p* = 0.061. An attempt to stratify on both sex and larval diet resulted in small strata and non-informative errors as expected (see Gordon, 2021). We therefore constructed a full model for data sets of 1) both sexes together, 2) females only, and 3) males only, and evaluated under each full model a full complement of sub-models using the dredge() function in the R Package MuMIn (Bartoń, 2022). The following models were analyzed:

1. Age_All_data_ ∼ Larval_Adult * strata(Sex) + Larval_Adult *Cage
2. Age_Females_ ∼ Larval_Adult* Cage
3. Age_Males_ ∼ Larval_Adult*Cage

We selected models based on model ranking using the Akaike information criteria for smaller samples (delta AICc and associated Akaike weights). Where models tied within 2 points of Λ1AICc, we summarized averaged competing models using the model.avg()in function MuMIn. Whenever models were averaged, we report on conditional averages only (i.e. the set of models actually averaged, not the full averaged model).

#### Fecundity and fitness

We performed several analyses on fecundity data. First, for each larval-adult diet regime, we calculated and visualized 1) lifetime group fecundity and 2) per female fecundity. Next, we calculated age specific fecundity from daily (i.e., 3 hours laying period) per female fecundity. We fit a linear model: Eggs ∼ Larval_Adult*Age + Larval_Adult*Cage and used dredge() to evaluate all possible models.

Next, we searched for patterns of sustained high and low egg periods across a time series using the changepoint R package [43]. Then, we compared fecundity patterns with either lifespan quantiles (25^th^, 50^th^, and 75^th^), or traditional mortality categories: early (10%), median (50%) and maximum lifespan (10%). Lastly, we calculated a proxy of fitness for females in each of the four treatments (L_C__A_C_, L_C__A_DR_, L_DR__A_C_ and L_DR__A_DR_) following the reasoning and assumptions in Matthews et al, (2021). We enumerated deaths and eggs 3x a week, thus we had a mix of two-day and three-day intervals. We thus computed age-specific survival as the mid-interval survivorship for individuals from age *x* to *x* + *n* days (*_n_L_x_*). We calculated age-specific fecundity (*m_x_*) at each interval as number of eggs in the interval (*b_x_*) divided by the number of females surviving, *k_x_* (i.e., *m_x =_ b_x_*/*k_x_*). We then used *_n_L_x_* and *m_x_* as input to construct Leslie matrices with assumptions as in [44] for each diet treatment. In the matrix, fecundity values form the top row while corresponding survival values line the diagonal, allowing to calculate the asymptotic population growth rate (lambda). Lambda is mathematically the dominant eigen value of the projection matrix and can be interpreted as a measure of fitness, or reproductive value [45] - the average fecundity of an individual expected at a given age as a measure of age-specific fitness. We used the function eigen.analysis() in the R package demogR [46] to calculate λ. With each reproductive value, we also extracted rho – a measure of the rate by which the population is expected to converge asymptotically to a stable distribution with a rate at least as fast as log(rho). Scripts and project notes to reproduce this analysis may be accessed at https://github.com/nochet/early_experience.

## Results

We analyzed patterns of survival and fecundity in eight populations of fruit flies reared in two diet conditions as larvae and split to each of the same diets as adults in two replicates, N > 300 individuals per replicate (Fig. 1). We calculated adult survival probability, per female fecundity, and a measure of fitness derived from combining lifespan and fecundity values at each sampling date. Our goal was to assess the effect of switched larval-adult diet regimes on the trajectories of these traits. Therefore, we focus on interaction effects of C_Larvae_ x C_Adult_, C_Larvae_ x DR_Adult_, DR_Larvae_ x C_Adult_ and DR_Larvae_ x DR_Adult_ diet treatment combinations (i.e. L_C__A_C_, L_C__A_DR_, L_DR__A_C_ and L_DR__A_DR_) while controlling for sex and cage (Table 1). The C diet is the *ad-libitum* feeding level while DR is protein-restricted. Phenotypes were recorded three times per week. Fecundities were observed for 3-hour periods just before lifespan recording. Overall, we observed four major patterns in survival: 1) survival is generally regime-dependent, 2) complex sex effects, 3) overall benefits of larval DR and 4) greater survival differences in pre- and post-median life phases (especially in males). Fecundity was overall 1) higher when larval diet was DR, but the timing of egg laying was advanced within the pre-median phase in A_C_’ treatments and delayed in ‘A_DR_’ flies (Table 1). In addition, substantial fecundities were observed for older post-median flies (> 50 days) in most treatments, and 5).

**Table 1:** Summary of survival and per female fecundity in each larval-adult treatment groups.

| Regime | Survival (%) | All |  |  | Female |  |  |  | Male |  |  |
| --- | --- | --- | --- | --- | --- | --- | --- | --- | --- | --- | --- |
|  |  | N | N<sub>ev</sub> | Days (CI) | N | N<sub>ev</sub> | Days (CI) | Eggs±se* | N | N<sub>ev</sub> | Days (CI) |
| C_C | 90 | 674 | 638 | 10(10-12) | 426 | 407 | 12(10-15) | 6.5±1.6 | 248 | 231 | 8(8-12) |
|  | 50 |  |  | 31(29-33) |  |  | 26(26-29) | 3.9±0.6 |  |  | 38(36-40) |
|  | 10 |  |  | 61(57-66) |  |  | 57(57-61) | 1.7±0.4 |  |  | 68(64-78) |
| C_DR | 90 | 721 | 676 | 12(10-15) | 393 | 370 | 15(10-17) | 6.8±1.1 | 328 | 306 | 10(8-19) |
|  | 50 |  |  | 38(36-40) |  |  | 36(36-40) | 8.5±2.1 |  |  | 38(36-40) |
|  | 10 |  |  | 71(68-78) |  |  | 64(61-68) | 1.0±0.2 |  |  | 87(85-92) |
| DR_C | 90 | 937 | 856 | 12(10-15) | 625 | 585 | 12(12-15) | 12.1±2.6 | 312 | 271 | 10(8-17) |
|  | 50 |  |  | 36(33-36) |  |  | 31(29-33) | 5.1±1.1 |  |  | 38(38-40) |
|  | 10 |  |  | 68(64-71) |  |  | 64(59-66) | 1.0±0.2 |  |  | 78(71-78) |
| DR_DR | 90 | 800 | 742 | 12(12-15) | 510 | 481 | 12(12-15) | 6.8±0.6 | 290 | 261 | 12(10-19) |
|  | 50 |  |  | 38(36-40) |  |  | 36(33-38) | 12.6±2.4 |  |  | 43(40-47) |
|  | 10 |  |  | 68(66-73) |  |  | 64(59-66) | 1.2±0.2 |  |  | 87(78-92) |

N is the number of individuals in a treatment, N<sub>ev</sub> is event count (i.e., deaths), Survival is the number of days for that treatment group to reach 90%, 50% and 10% survival (i.e., 10%, 50%, 90% mortality), CI, confidence intervals around survival estimates, se, standard error of mean eggs. \*Mean number of eggs per female per 3 hour-laying period.

### Survival patterns depend on diet regime

The effect of larval-adult diet combinations on the pattern of survival in mixed-sex groups was subtle but differed significantly (log-rank, df = 3, χ^2^ = 48.2, *p* = 2e^-10^, Fig. S1). Survival trajectories diverged from about 70% survival and indicate a complex pattern especially after median survival (i.e. from about 38 days onward). Survival was lowest in L_C__A_C_ flies overall with a median of 31 days (Table 1), suggesting that constant *ad lib* feeding from larval to adult age limits survival (Table 1, Fig. S1). Compared to L_C__A_DR_ and L_DR__A_DR_, the L_DR__A_C_ curve showed a steeper mortality patten after median survival, like that seen in L_C__A_C_ flies (Fig. S1), suggesting that developmental dietary restriction followed by *ad libitum* feeding is a less optimal sequence for longevity. We note however that this sequence improves survival in the early half of lifespan compared to L_C__A_C_. The fact that L_C__A_DR_ and L_DR__A_DR_ incurred the least mortality rates might suggest that adult diet, especially if it is DR, has a larger contribution to longevity.

### Survival was significantly sex-dependent in diet regimes

As expected, exploratory analysis showed sex in addition to diet regime effects (Table 1). To get a sense of the extent of sex differences, we visualized survival for L_C_ and L_DR_ for sexes separately (Fig. 2).

**Figure 2:**
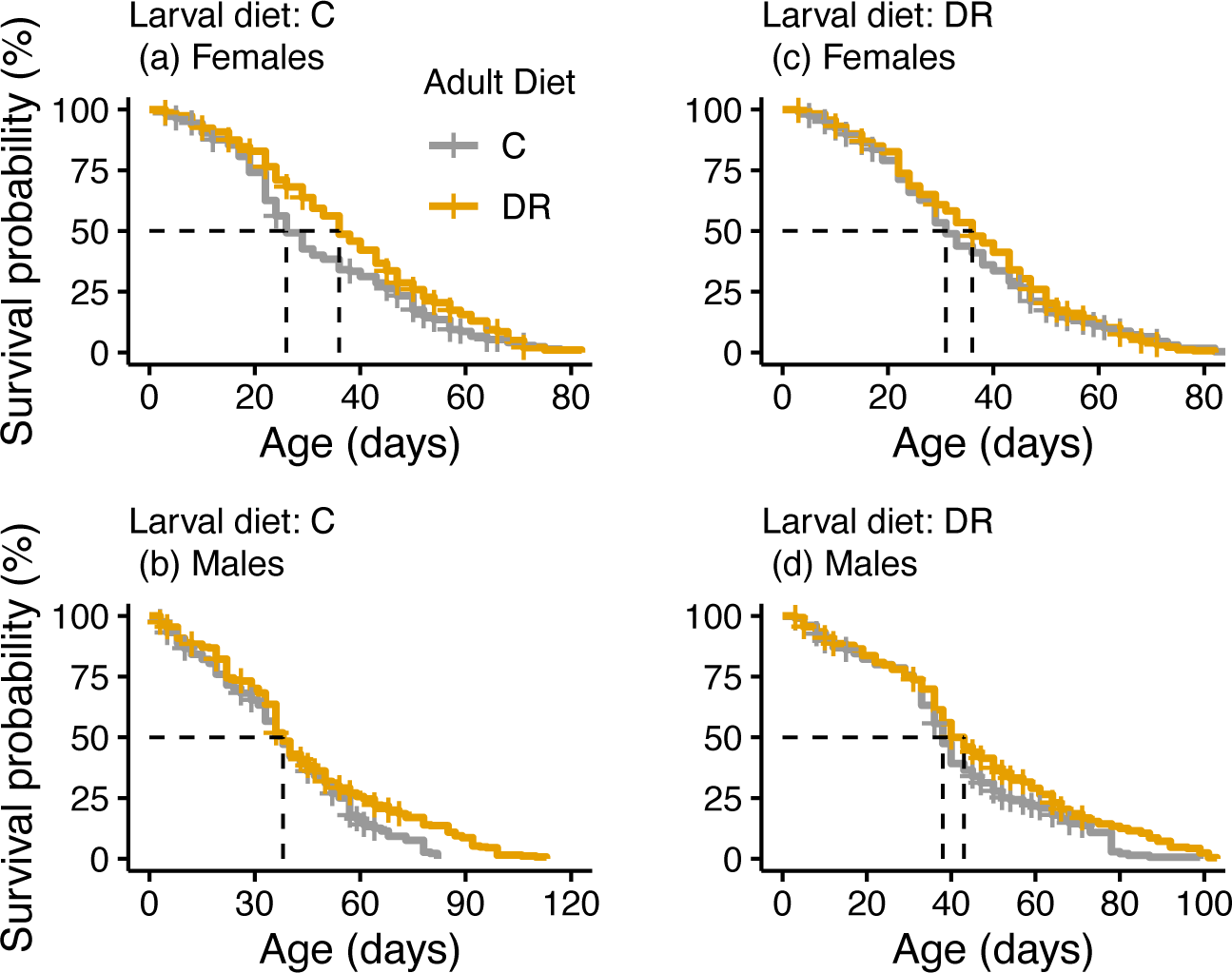
Probability of survival grouped by larval-adult diet regime and sex. Broken lines depict age (days) at median survival probability. Survival responded in an age-specific manner to treatment combinations, and differently between the sexes.

Shortest and longest median survival appeared in L_C__A_C_ females (26 days, CI: 26-29 days) and L_DR__A_DR_ males (43 days, CI:40-47 days), respectively (Fig. 2a&d, Table 1). To the contrary, no differences between L_C__A_C_, L_C__A_DR_ and L_DR__A_C_ in male median lifespan (38 days). While males reared on DR constantly (i.e., L_DR__A_DR_) had longest median lifespan (43 days), female L_DR__A_DR_ had a much lower median (36 days, CI: 33-38 days) which tied with L_C__A_DR_ females (36 days, CI: 36-40 days). When larval diet was DR, median lifespan is also consistently shorter for females (31 days for L_DR__A_C;_ 36 days L_DR__A_DR)_ than males (38 days L_DR__A_C;_ 43 days L_DR__A_DR_). Lastly, we further note that diet switching between larvae and adult stages resulted in generally intermediate median lifespans. The finding that matched larval-adult diets yielded the least and greatest medians is one we return to later. Overall, these results underscore complex response patterns to life stage-based diet variability.

#### Greater effects of diet in post-median lifespan and longevity

Lifespan was more variable among regimes in the post-median survival phase, after 36 days of age especially in males. Males lived variably longer in maximum lifespan (i.e. 10% survival): L_C__A_C_ 68 days, L_C__A_DR_ 87 days, L_DR__A_C_ 78 days, and L_DR__A_DR_ 87 days; mean 80 days. Females lived 16 fewer days on average in maximum lifespan, and remarkably similar across regimes L_C__A_C_ 57 days, L_C__A_DR_ 64 days, L_DR__A_C_ 64 days, and L_DR__A_DR_ 64 days (Table 1). L_C__A_C_ females with the least median lifespan, maintained the lowest maximum survival (57 days, CI: 57-61 days). We note that the L_C__A_DR_ males tying with L_DR__A_DR_ males at 87 days in maximum lifespan had a median like all treatments, suggesting that they reached high maximum lifespan by slowing their post-median mortality. Thus, the pattern of lifespan post-median differed from that pre-median, and the post-median trajectories contribute significantly to a complex response involving both diet regimes and sex. In general, there were greater gains in maximum lifespan for males than females in most regimes over the L_C__A_C_ regime.

#### Life phase-based survival response to larval-adult diet regimes

From results presented, 1) female performance in median and maximum lifespan was generally low, 2) treatments involving DR as adult diet performed better on average than those with C adult diet, particularly in the median lifespan, and 3) matched larval-adult diets were inconsistent outcomes, producing the shortest- and longest-lived groups. Rather, we observed a pattern that highlights sex and adult stage as interacting with diet regime. Fourthly and importantly, there is generally little variability lifespan around the median lifespan phase compared to the pre- and post-median phases. We examine the last two points further in the following sections. The rest of our analysis address the last two points using a modeling approach to identify 1) sources of variation for matched vs mismatched diet regime outcomes, and 2) identify phases of adult life associated with longer or shorter life in early, median, and maximum lifespan.

#### Variation in the risk of death in matched and mismatched diet regimes

To further understand the effect of larval-adult diet sequence on survival, we constructed a Cox model containing all major terms (i.e. Age ∼ L_A * s(S) + L_A*C) and systematically evaluated all sub-models for combined male and female data (see methods). Performed on all data we found a significant effect of cage, with several models tied in ΔAICc weights (Table 2), and no effects of diet in conditional averages. A marked effect of cage might in part arise from the fact that there were generally twice as female many events compared to male events except for L_C__A_DR_ (Table 1). To address this issue, and because exploratory analysis suggested role of sex effects, we repeated our model on each sex separately. For females, several models were tied within 2 AICc points. The third of these had one term, L_A with: coeff. -0.23, se 0.07 *p* <0.001; coeff. -0.13, se 0.06, *p* < 0.05; and coeff. -0.20, se 0.07, *p* <0.01, respectively for L_C__A_DR_, L_DR__A_C,_ and L_DR__A_DR_, relative to L_C__A_C_. For males, one model containing one term (L_A) was identified: N_ev_=1069, coeff. -0.31, se 0.09 *p* <0.001; coeff. -0.12, se 0.09, *p* = ns; and coeff. -0.36, se 0.09, *p* <0.001, respectively for L_C__A_DR_, L_DR__A_C,_ and L_DR__A_DR_, relative to L_C__A_C_. These results reveal a larger effect of diet regime in males than females. Since both mismatched L_C__A_DR_ and matched L_DR__A_DR_ are significantly different from matched L_C__A_C_, while the unmatched L_DR__A_C_ is not different from the matched L_C__A_C_, matched diet environments do not necessarily result in lower risk of death, at least in males. A similar, but abated pattern holds for females except that the L_DR__A_C_ regime was different from L_C__A_C_. These results highlight the potential negative effects on lifespan of switching from a rich larval diet (C) to a poor adult diet (DR). Detailed model results are presented in Table S1.

**Table 2:**
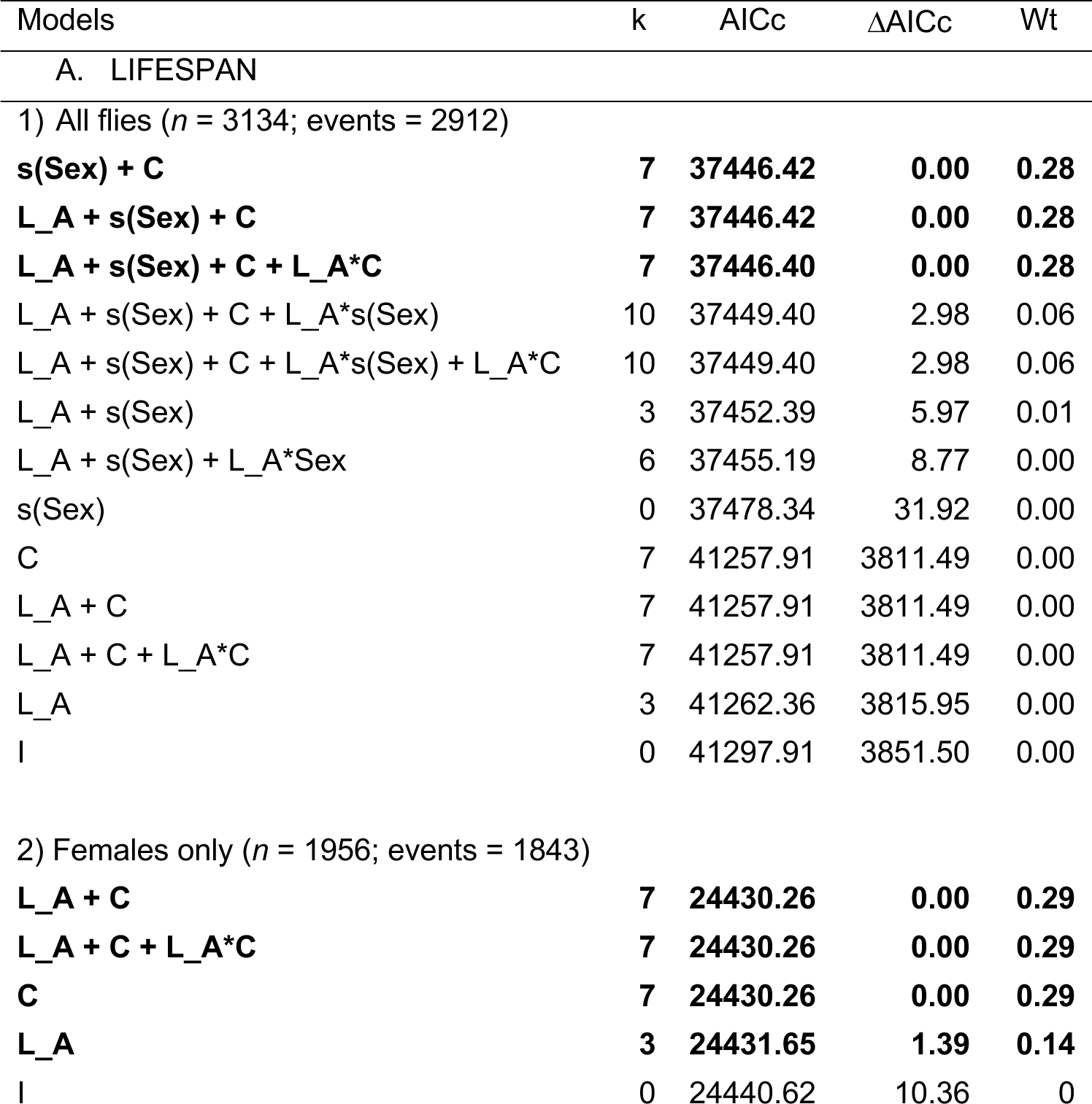
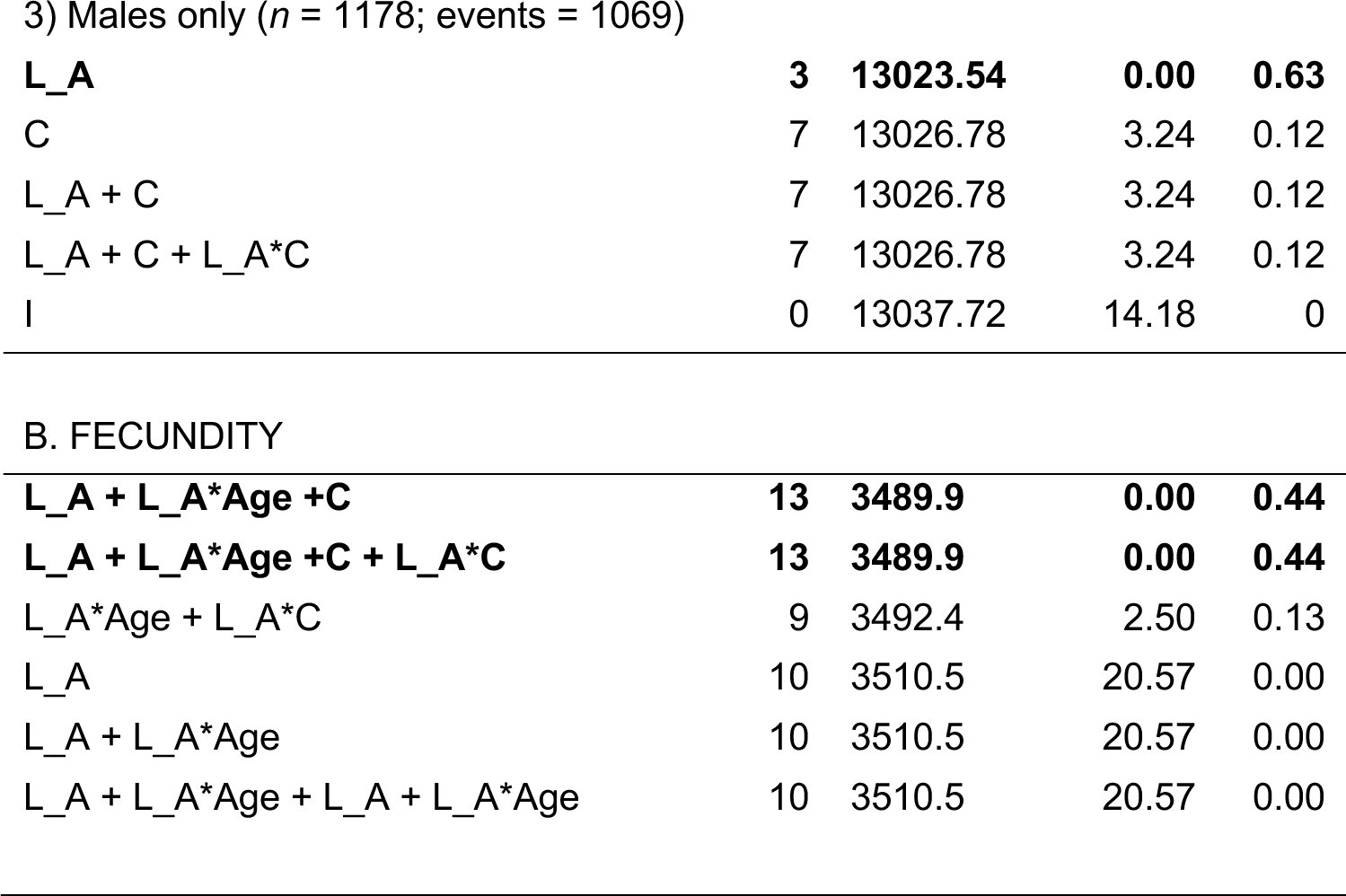
Selection of Cox models. The best modes are highlighted in bold face.

| Models | k | AICc | $\Delta AICc$ | Wt |
| --- | --- | --- | --- | --- |
| A. LIFESPAN |  |  |  |  |
| 1) All flies ($n = 3134$; events = 2912) | | | | |
| **s(Sex) + C** | **7** | **37446.42** | **0.00** | **0.28** |
| **L_A + s(Sex) + C** | **7** | **37446.42** | **0.00** | **0.28** |
| **L_A + s(Sex) + C + L_A\*C** | **7** | **37446.40** | **0.00** | **0.28** |
| L_A + s(Sex) + C + L_A*s(Sex) | 10 | 37449.40 | 2.98 | 0.06 |
| L_A + s(Sex) + C + L_A*s(Sex) + L_A*C | 10 | 37449.40 | 2.98 | 0.06 |
| L_A + s(Sex) | 3 | 37452.39 | 5.97 | 0.01 |
| L_A + s(Sex) + L_A*Sex | 6 | 37455.19 | 8.77 | 0.00 |
| s(Sex) | 0 | 37478.34 | 31.92 | 0.00 |
| C | 7 | 41257.91 | 3811.49 | 0.00 |
| L_A + C | 7 | 41257.91 | 3811.49 | 0.00 |
| L_A + C + L_A*C | 7 | 41257.91 | 3811.49 | 0.00 |
| L_A | 3 | 41262.36 | 3815.95 | 0.00 |
| I | 0 | 41297.91 | 3851.50 | 0.00 |
| 2) Females only ($n = 1956$; events = 1843) | | | | |
| **L_A + C** | **7** | **24430.26** | **0.00** | **0.29** |
| **L_A + C + L_A\*C** | **7** | **24430.26** | **0.00** | **0.29** |
| **C** | **7** | **24430.26** | **0.00** | **0.29** |
| **L_A** | **3** | **24431.65** | **1.39** | **0.14** |
| I | 0 | 24440.62 | 10.36 | 0 |
| 3) Males only ( $n = 1178$ ; events = 1069) |  |  |  |  |
| **L_A** | **3** | **13023.54** | **0.00** | **0.63** |
| C | 7 | 13026.78 | 3.24 | 0.12 |
| L_A + C | 7 | 13026.78 | 3.24 | 0.12 |
| L_A + C + L_A*C | 7 | 13026.78 | 3.24 | 0.12 |
| I | 0 | 13037.72 | 14.18 | 0 |
| B. FECUNDITY |  |  |  |  |
| **L_A + L_A\*Age +C** | **13** | **3489.9** | **0.00** | **0.44** |
| **L_A + L_A\*Age +C + L_A\*C** | **13** | **3489.9** | **0.00** | **0.44** |
| L_A*Age + L_A*C | 9 | 3492.4 | 2.50 | 0.13 |
| L_A | 10 | 3510.5 | 20.57 | 0.00 |
| L_A + L_A*Age | 10 | 3510.5 | 20.57 | 0.00 |
| L_A + L_A*Age + L_A + L_A*Age | 10 | 3510.5 | 20.57 | 0.00 |

#### Protein-restricted larval diet improved lifetime fecundity

We recorded the number of eggs laid on media in each cage in a 3-hour period, three time per week over lifetime (N = 2 cages per diet regime, 8 cages in total). These are same cages in which lifespan was measured, providing opportunity to compare fecundity and lifespan performance in each diet condition. Lifetime fecundity in diet groups was varied: L_C__A_C_ = 3664, L_C__A_DR_ = 3432, L_DR__A_C_ = 9795, and L_DR__A_DR_ = 5533 total eggs counted. In a 3-hour laying period an L_DR__A_C_ female on average laid 1.2 eggs, compared to <0.5 eggs in the rest (Fig. 3). These counts suggest that moving to a richer diet as adults favors higher fecundity compared to all other combinations. Results further suggest that both matched and mismatched diet conditions can have vastly different responses by adjusting timing and duration of reproductive activity. General patterns of fecundity are visualized in Figures 3 and S3.

**Figure 3:**
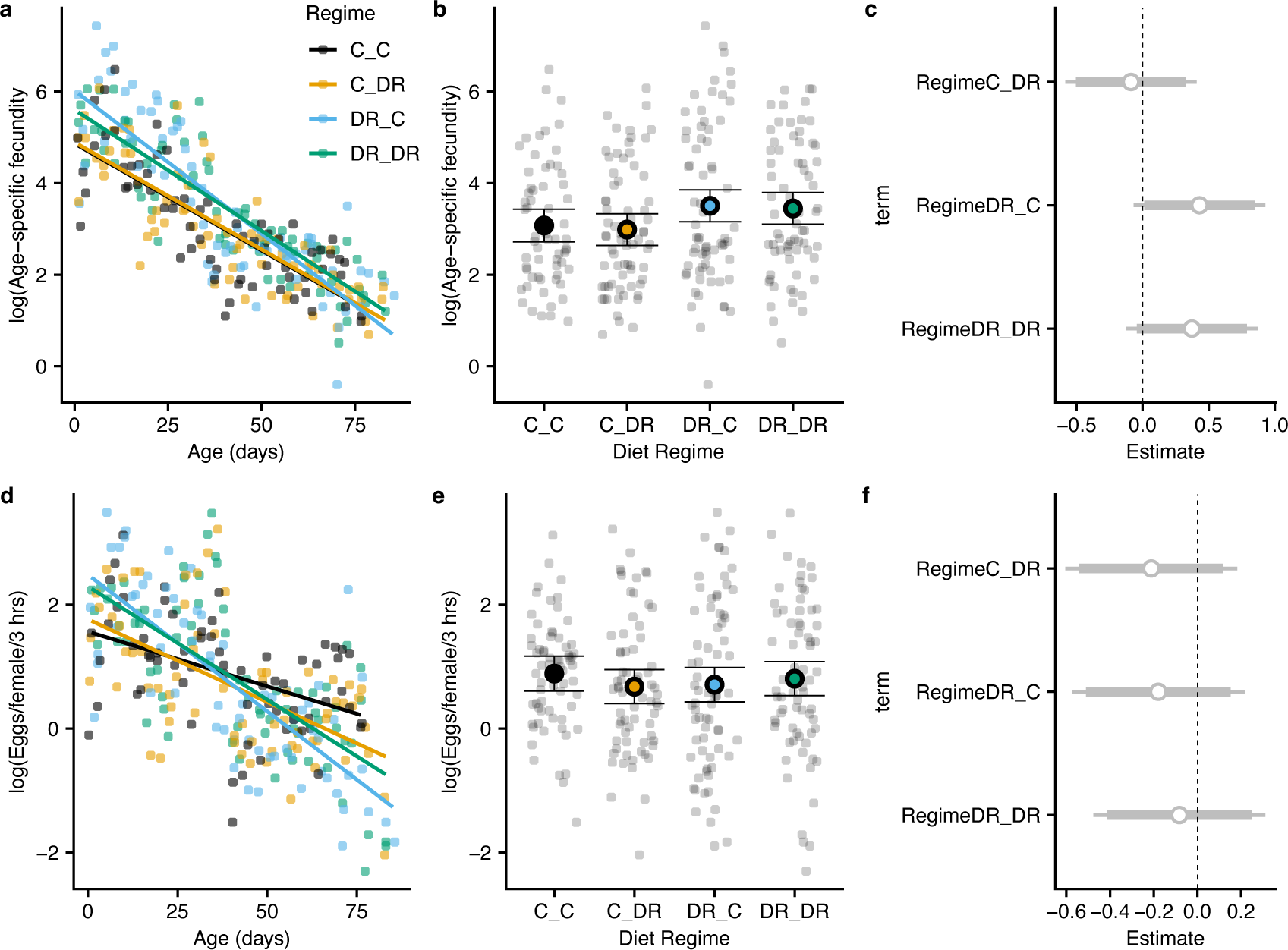
Pattern of egg output in each treatment of larval-adult diet regime: **a-c** – Mean age-specific fecundity (i.e. mean number of eggs laid in a 3-hour period in each diet regime over lifespan is plotted. Lines are linear models of those diets; **d-f** – same data in a-c calculated per female; **c&f** – are estimated regime effects calculated from regressing numbers of eggs on regime, ether in group fecundity **c** or per female fecundity **f**.

#### ‘Low to high’ diet transition favors higher age-specific female fecundity

We found significant differences in age-specific fecundity for two comparisons (L_DR__A_C_ vs L_C__A_C_, *p* <0.05 and L_DR__A_C_ vs L_C__A_DR_, *p* <0.05, pairwise t-test). We note that L_DR__A_C_ was 5 days lower in median lifespan compared with both L_C__A_DR_ and L_DR__A_DR_ but tied with L_C__A_DR_ in female maximum lifespan at 64 days. We found a significant interaction effect of age with the L_DR__A_C_ regime (linear model est. -4.1, z 4.02, *p* <0.001). Finally, as expected, increasing age had a negative effect on age-specific fecundity overall (est. -2.5, z 3.3, *p* <0.001). Unlike lifespan, cage effects were limited to both cages of only the L_DR__A_C_ treatment. Model results are reported in Table S1.

#### Diet regime influenced timing and duration of peak fecundity

The age-specific pattern in fecundity revealed peak laying periods that were variably early or late but seemed unique to each regime, suggesting stages or phases of egg laying. Although most eggs were expectedly produced earlier in life (15 - 38 days *po*), obvious patterns of prolonged dips and spikes were observed (Figs. S3). To gain further insight into this pattern, we looked for significant change points in the patterns of egg laying across lifespan in each treatment. We identified a pattern in which both regimes starting with larval DR diet showed a longer initial period of sustained oviposition compared to treatments starting out with the C diet (Fig. 4) although the specific character of these episodes differs. Further, although fecundity plumates in all treatments after about 50 days, there remains substantial egg laying, and a few cases, some low-grade spikes in old females (Fig. S3).

**Figure 4:**
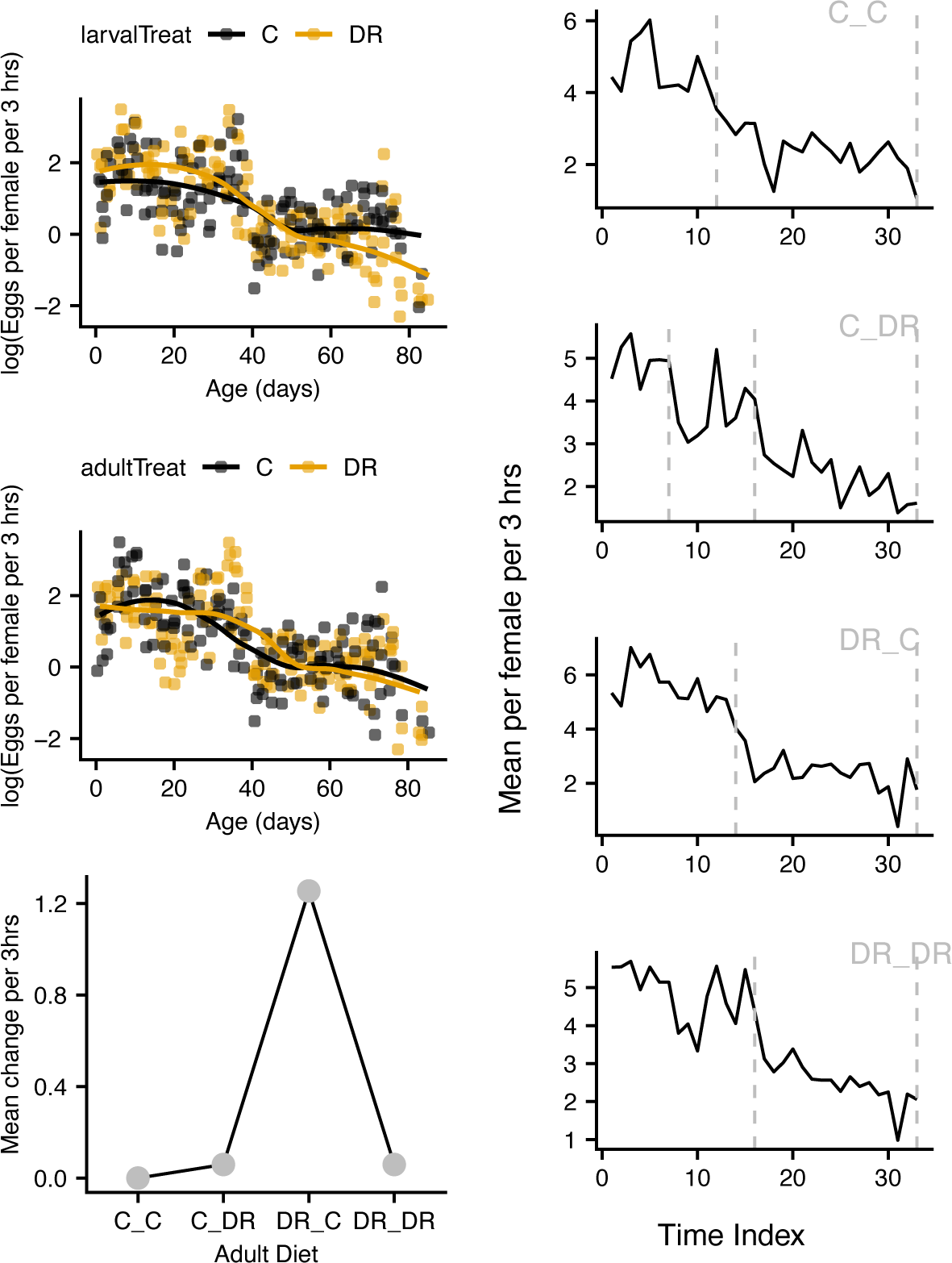
Average number of eggs per female per day calculated from 3-hour laying periods sampled 3 times a week over lifetime. *Top left* – grouped by larval treatment C or DR. *Mid-left* – grouped by adult treatment C or DR. *Bottom left* – Mean difference in egg number per 3-hour laying period in each regime. *Right* – Number of detected change points in egg-laying and duration of these periods.

### Female fecundity in the context of survival patterns

In this study, lifespan was lowest in a matched larval-adult diet (L_C__A_C_) and tied at highest in matched (L_DR__A_DR_) and mismatched L_C__A_DR_ in both median and maximum lifespan. Similarly, fecundity was lowest in matched L_C__A_C_ treatment and highest in a mismatched L_DR__A_C_. A switch from C to DR favors lifespan while that from DR to C favors reproduction.

To confirm this observation, we combined age-specific lifespan and fecundity values for each treatment into a fitness proxy measure using life tables and Leslie matrix calculations (see methods) and obtained a reproductive value (ƛ) and its associated damping ratio (rho) in each condition: L_C__A_C_, 2.6, 3.7, 24.0; L_C__A_DR_, 3.5, 3.7; L_DR__A_C_, 3.5, 2.8; and L_DR__A_DR_, 3.2, 8.9.

Figure 5 presents a comparison of age-specific reproductive values (i.e. the average individual female fecundity to be expected at that age) between treatments. Trends in the plots are typically different for each of matched or unmatched pairs. L_C__A_C_ and L_DR__A_C_ decline almost monotonously before leveling out in old age. In L_C__A_DR_ and L_DR__A_DR_ peak fecundity shifts to the right along the time axis representing a delay in their peak fitness performance. These trends generally hold when fecundity data is analyzed on its own (Fig. S4) or in lifespan quantiles (Fig. S5). Our results do not appear to support the silver spoon and environmental matching hypotheses.

**Figure 5:**
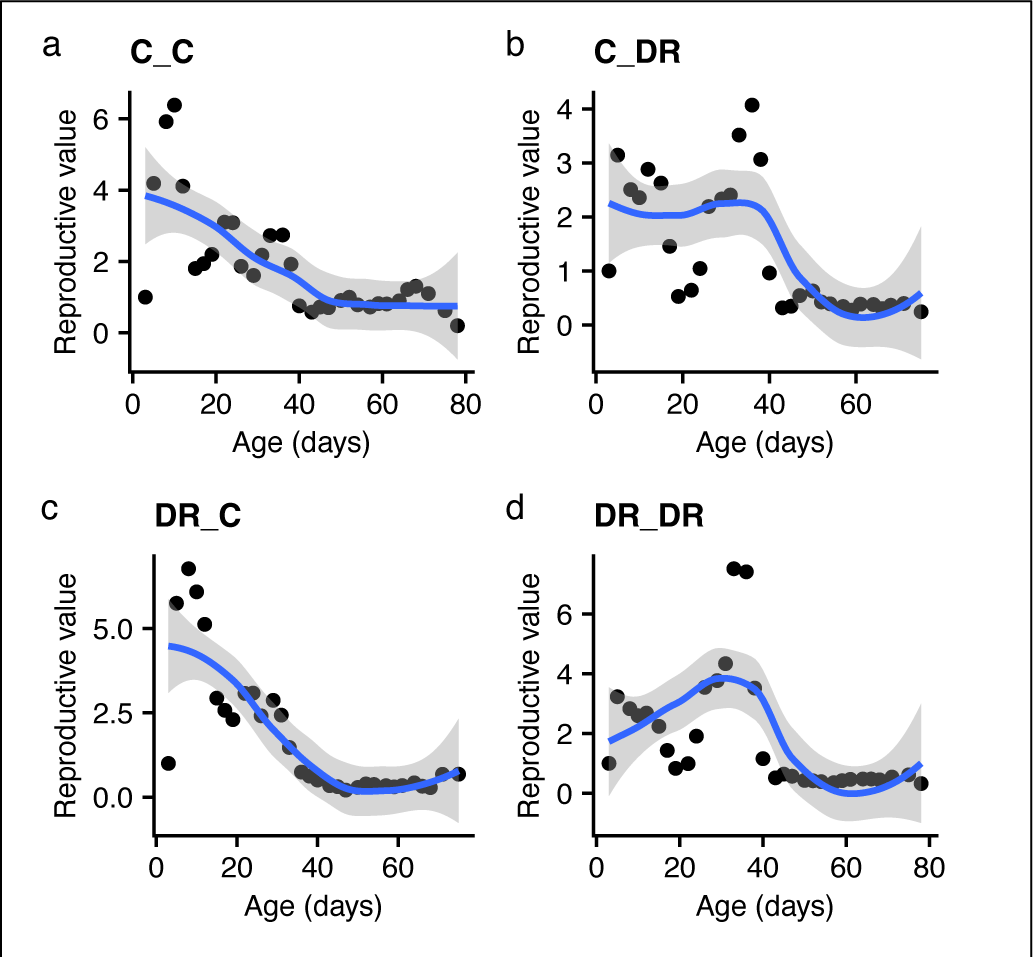
Age-specific fecundity patterns in each of the four diet combinations. A reproductive value is the expected per 3-hour fecundity of a female calculated from lifespanfecundity Leslie matrices. The dots represent mean per female fecundity at each sampling time over the lifespan.

## Discussion

We studied fitness effects of developmental-adult nutrient availability in an outbred population generated from 835 lines of a multiparent *D. melanogaster* population. Fitness in this study is a proxy obtained from full lifespans and partial fecundity measured over full lifespan in each treatment (see methods). We address the question “what is the developmental x adult diet interaction effect of low/high protein on adult reproduction and lifespan in a high genetic diversity multiparent population?” We anchored our expectations on two competing hypotheses of developmental plasticity as reviewed in Monaghan [6]. The environmental matching hypothesis emphasizes the constancy of environmental condition across life stages as critical for maximizing fitness. This means generally, that good or poor conditions if maintained that way across lifespan will favor higher fitness for both, but those who experienced better early condition will be higher than those with poor conditions (i.e. parallel norms of reaction). On the other hand, the silver spoon hypothesis assumes greater fitness outcomes of better developmental conditions, regardless of adult conditions (although those with good conditions in both phases will be higher than those with good early but bad adult conditions). Thus, organisms that developed in better conditions are predicted to be primed for higher fitness. We tested fitness outcomes in the following conditions: L_C__A_C_ (low start, low adult), L_C__A_DR_ (low start, high adult), L_DR__A_C_ (high start, low adult) and L_DR__A_DR_ (high start, high adult).

We do not find evidence for either model, but observed several major patterns: 1) survival was age-specific in all diet regimes, temporary differed across regimes, and differed between sexes, 2) major survival differences were seen in post-median life phase, especially for males (although this effect is, to a lesser degree, apparent particularly in females receiving the richer control diet), 3 survival was both longest and shortest in matched larval-adult diets; fecundity was higher when early diet was DR, 4) matched and mismatched involving DR as larval diet showed delayed and advanced laying periods, respectively.

### Diet matching or mismatching each yielded high and low fitness

Under the matching hypothesis, the L_C__A_C_ treatment should lead to greater fitness compared to mismatched treatments L_C__A_DR_ and L_DR__A_C_ (i.e., more offspring). The matched L_DR__A_DR_ should lead to lowest fitness performance compared to L_C__A_C_, but better than in switched combinations (L_C__A_DR_ and L_DR__A_C_). We observed highest fecundity in a mismatched regime, L_DR__A_C_. Further, regimes starting out low (L_DR__A_C_ and L_DR__A_DR_) had significantly larger lifetime egg output. Similarly, under silver spoon, better larval diets (L_C__A_C_ and L_C__A_DR_) should show best fitness performance (i.e., the early diet matters more). Both achieved comparatively poor fitness outcomes with the lowest reproductive value of 2.6 in L_C__A_C_.

Given that these hypotheses have been confirmed in some studies, we speculate on the role of plasticity in our population, especially that the level of protein restriction (i.e. 50%) is fairly easy to adjust to. A study that considered adaptive responses to nutritional challenges in the context of experimental evolution reported faster development and decreased adult weight as response to a lower calorie developmental diet (May et al, 2019). Morphological changes in say, ovariole size or number observed in some studies (e.g. Deas et al. [4], would be unlikely in our case in one generation under relatively low larvae and adult density we reared our flies in. Our findings are reflected in a study of the African clawed frog similar in design to this study [47]. Manipulating dietary protein, they measured hormonal effects on vocal ability of males, and confirmed neither hypothesis.

### Restricted developmental diet improved fitness

Regardless of combination, crossed or matched, early DR diet was associated with higher fecundity. This was true generally for lifespan also, except for the 5 days reduction in median lifespan in the L_DR__A_C_, which also showed nearly twice as many eggs as other regimes. This regime also advanced fecundity in producing an early peak (Fig. 4). We therefore interpret lower median survival in L_DR__A_C_ as arising from a steeper survival-fecundity trade-off. But why would a relatively low protein diet favor increased fitness? It is conceivable that significant resources are allocated to locomotor activity, and this becomes more important in longitudinal studies like ours. A study in *D. melanogaster* in which in addition to fecundity and lifespan, measured activity level also in a series of protein restricted diets (30-90% protein), concluded that chronic protein restriction in all diets from eclosion did not induce malnutrition [48]. Thus, apart from the key lifespan-reproduction trade-off, high fecundity in DR treatments might relate to improved locomotor function and generally a slower pace in functional senescence [49]. With slower declines in activity, DR flies may engage in more reproductive activity longer compared with those on high protein diet.

As for why low protein in development should result in high fecundity, we speculate that deficits in development were potentially compensated for by a significant delay in peak laying as adults (i.e. the L_DR__A_C_ regime). Where this compensation is limited, a cost is paid in median lifespan (i.e. 5 days lower in LDR_A_C_). In the bank vole (*Myodes glareolus*), Schroderus et al, [50] could not explain evident phenotypic trade-offs in offspring number vs size by a genetic trade-off, but rather by negative correlations in environmental effects. In that study, some genetic correlations estimated were positive leading them to conclude that genetic variation in those traits may not always be antagonistic in mammals. Our results shine a spotlight on potential non-genetic environmental effects.

### Other interesting patterns

We observed significant and sex-based differences in survival and reproductive output in post-median life phase. Post-median lifespan was long in both sexes but pronounced in males. We further have reported on stage-rather than age patterns in egg production (change point analysis). Simulation studies [51], suggest that exposure to different environmental perturbations can trigger different age-specific demographic rates which might lead to divergent age-specific vs stage-specific patterns especially in the presence of strong survival-fecundity trade-offs (the case of the L_DR__A_C_ regime). Stage-associated early-late diet effects on fitness might suggest the presence of different underlying response mechanisms recruited in diet combinations.

Males generally survived longer, and independent of diet treatment in median and especially maximum lifespan than females in this study. Several explanations might account for this result. From the energetics point of view, sexual size dimorphism can lead to differences in sensitivity to environmental conditions during the growth period between the sexes [5,52,53]. This dimorphism may be due to different energetic requirements for males and females. The heavier sex is likely to be more vulnerable to low food availability. While dimorphism may not be apparent at the larval stage, low growth conditions might predispose larger individuals in adulthood (typical in females) to greater risk of death. Further, *D. melanogaster* exhibit sexual dimorphism in foraging choice [52,54]. When presented with choice, Davies et al [52] found that adult females opted for the same protein-to-carbohydrate (P:C) ratio regardless of their developmental diet. Males on the other hand preferred a diet that compensated for deficiencies – if protein was low during development, males increased protein consumption. If no choice is provided (as in this study), adult lifespan increased as P:C ratio decreased, independent of developmental diet.

Alternatively, high male density can lead to heavier males on average. In the butterfly *Bicyclus anynana*, [55] found that high male density favored production of larger body mass and storage, which increased male survival in times of intense intraspecific competition through decreased courtship activity and sperm number. In this study, we exclude effects of density. Both, during collection of eggs for experimental populations and experimental cages themselves were stocked at about a third of the maximum density (1.e. about 600 - 900 flies in cages designed >3000 flies).

In this study, we did not control the ratio of males to females. While it is conceivable that male-male competition for mating can be energetically costly and result in reduced male lifespan overall, the mere presence of large numbers of males in our cages is enough to stress females and raise their mortality risk. Indeed, [56] analyzed climbing rate and starvation resistance in *D. melanogaster* and found that mere exposure of females to males, and mating accelerated female functional aging, leading to declines in both traits, and this effect was independent of changes in body mass, fat storage, and sex peptides.

Our study used outbred fruit flies likely much higher in the level of standing genetic diversity. Many fruit fly studies are performed in inbred lines, and typically a few lines. Here, we focused on the combined effect of larval-adult diet. Many studies we have reviewed above explored effects of a developmental diet, or additive effects of developmental and adult diets. Our key findings do not support silver spoon and environmental matching hypotheses, as others have also found. Our results may reflect this particular population and the specific question we set out to answer. Future studies should invest in uncovering underlying genetic mechanisms for these striking patterns.

### Conclusions

Theory of developmental plasticity is a traditional framework for explaining patterns of fitness differences in populations experiencing different conditions across ontogeny. We studied the joint effect of larval and adult diet differing only in the quantity of protein. Our test population was outbred from a large number of inbred lines and was initially founded from a global set of eight founder lines. Our results are not consistent with the predictions of either the silver spoon or environmental matching hypotheses of developmental plasticity. Results also suggest increased fitness in protein restricted treatments, and differences in the timing of fecundity allocation triggered by diet interactions. We attribute these results to high standing genetic diversity present in this multiparent resource. In addition, an outbred population might present allelic options for responding to environmental change. Studies aiming to understanding adult trait expression should account for ontogeny-wide organismal conditions.

## Acknowledgements

Anna Perinchery-Herman constructed the lighting studio box we used for image capture.

## Authors’ contributions

EN conceived the idea; EN, EGK and AJ designed the study; AJ and EN set up the experiment; AJ collected lifespan and egg image data; AJ, EN, and DD extracted counts from images; AJ and EN performed data analysis; EN drafted the manuscript; AJ, DD, AND EGK provided comments on the manuscript. All authors approved the manuscript for publication.

## Competing interests

We declare we have no competing interests.

## Funding

This work was supported by NIH grant no. R01 GM117135 to EGK, and the University of Missouri startup funds to EN.

## Notes

https://zenodo.org/records/10445786

https://github.com/nochet/early_experience

### Competing Interest Statement

The authors have declared no competing interest.

