## Supplemental File 1 for "Larval protein restriction interacts with adult diet to increase fitness in an outbred multiparent population"

Supplementary materials

Female fecundity trends in age classes defined in a survival model (age classes here).

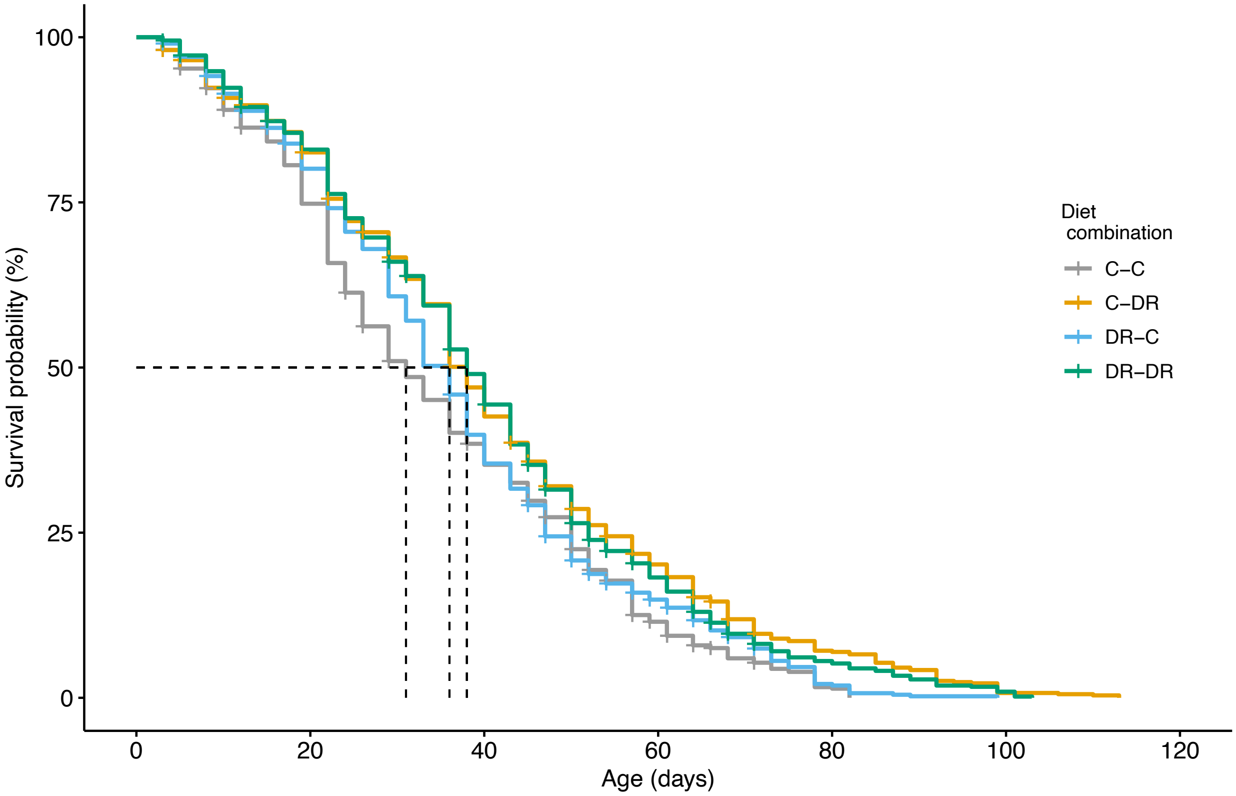

Figure S1: Kaplan-Meier survival probabilities for each of the larva-adult diet combination for both sexes ($X^2 = 47.1$, df = 3, $P$ <3e-10). Shortest and longest median survival were observed in C-C and DR-DR.

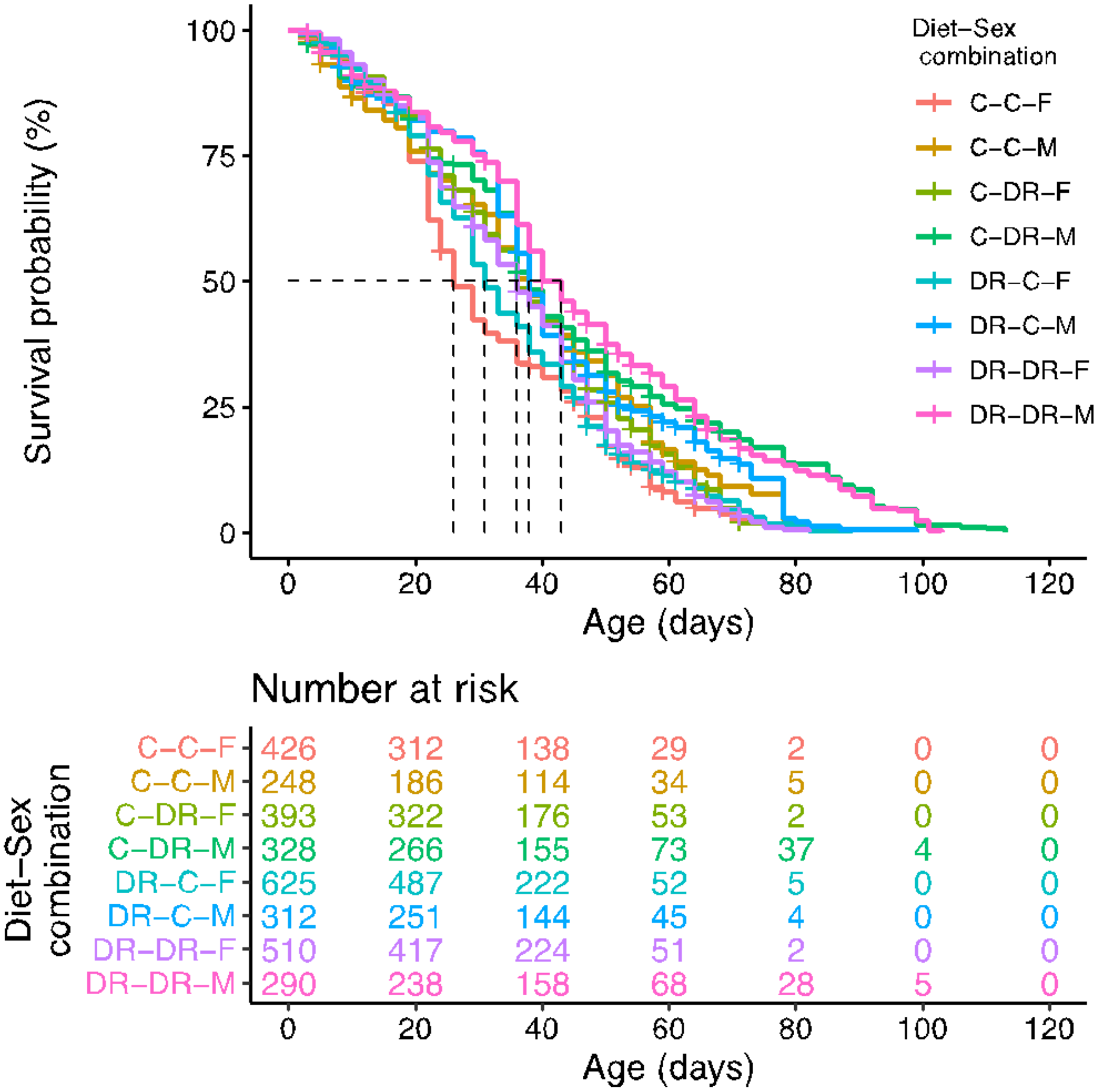

Figure S2: Kaplan-Meier survival probabilities for each of the larva-adult diet combination for each sex ($X^2$ = 148, df = 7, *P* <2e-16). Shortest and longest median survival were observed in C-C-F and DR-DR-M. Interestingly, sex reciprocals of these were intermediate in the band of curves. Males in the C-DR and DR-DR combinations showed >10 % at 80 days of age, indicating slower aging in these diets.

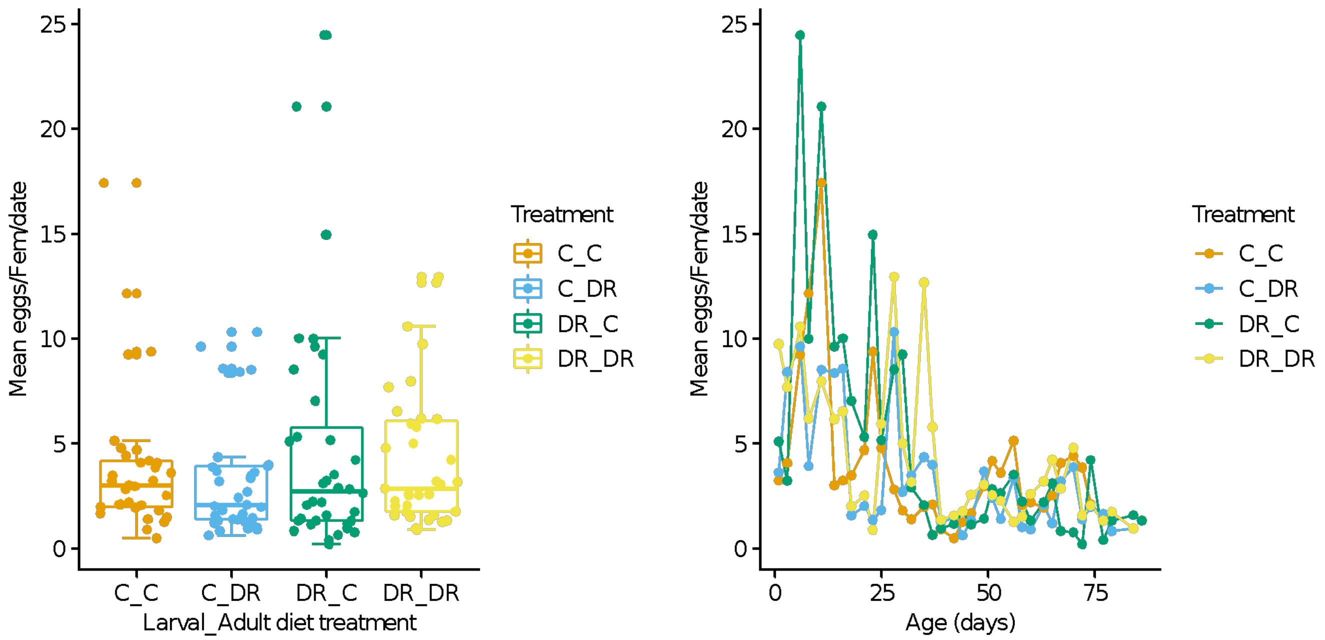

**Figure S3:** Average number of eggs per female per day calculated from 3-hour laying periods sampled 3 times a week over lifetime. **A.** Pooled over lifetime: points are means for each sampling date. **B.** A time-series showing means for each collecting date. C_C, larval C & adult C diet; C_DR, larval C & adult DR; DR_C, larval DR & adult C; DR_DR, on DR diet.

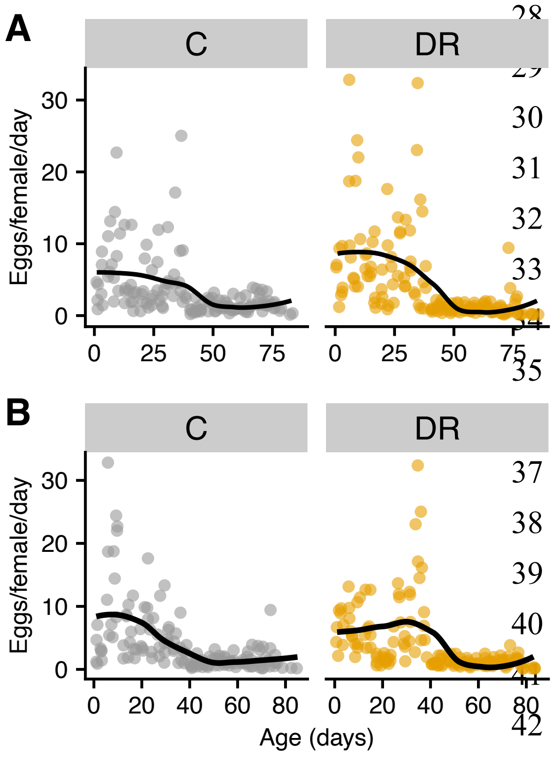

**Figure S4:** Fecundity plotted separately for each larval diet: **A** left, larval C and adult C; **A** right, larval DR and adult DR; **B** left, adult C and early DR; **B** right, larval DR, adult C.

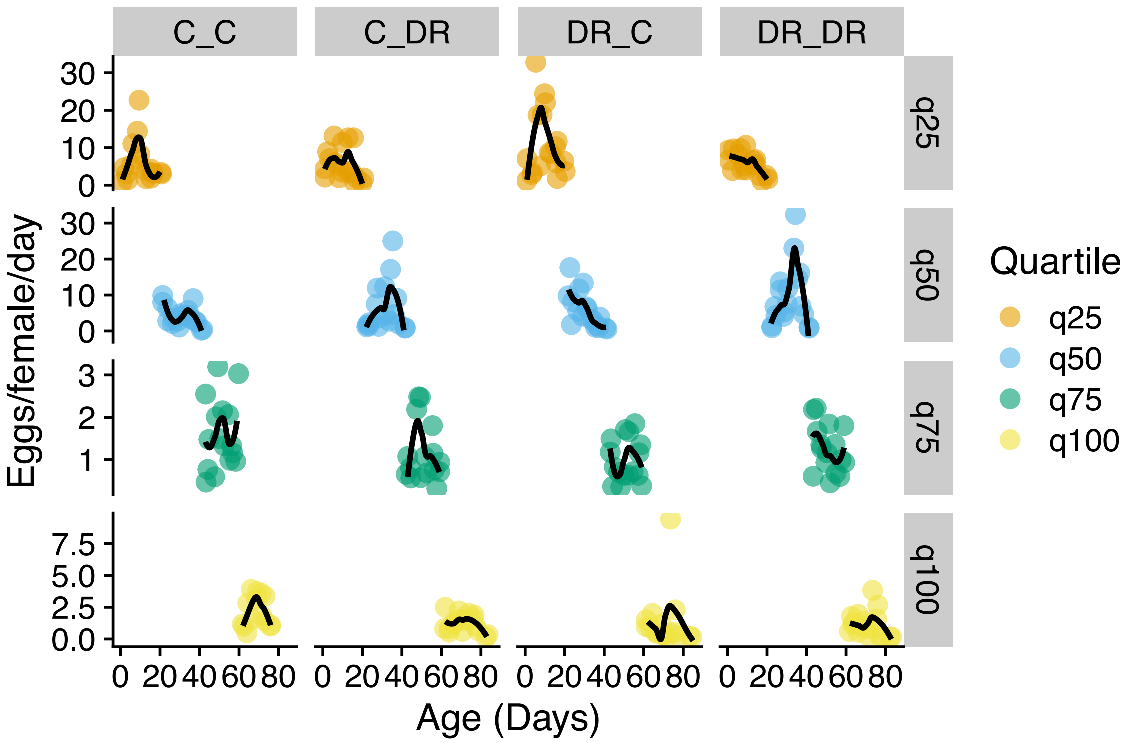

**Figure S5**: Fecundity patterns estimated in each quantile of fruit fly lifespan in each diet regime.

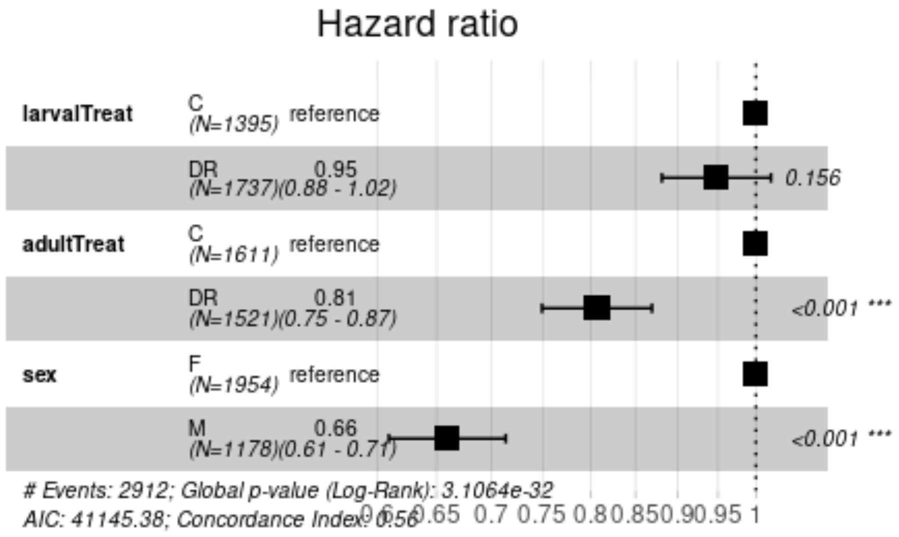

**Figure S6**: The risk of death in each diet regime obtained by fitting a cox model. Note, in this model sex is not fit as strata(sex) reported in the main text. This model overestimated the risk of death as sex violated the proportionality assumption of the cox model. We show it here only to provide comparison with corrected hazards reported.

**Table 1**: Detailed model results. For averaged models, we interpret conditional averages.

| | Estimate | Std. error | z | Pr(>\|z\|) | |
|---|---|---|---|---|---|
| **Lifespan all flies: Model-averaged coefficients** | | | | | |
| Full average | | | | | |
| cageIDC_C2 | 0.01647 | 0.07973 | 0.207 | 0.8364 | |
| cageIDC_DR1 | -0.15564 | 0.08730 | 1.783 | 0.0746 | . |
| cageIDC_DR2 | -0.33291 | 0.07691 | 4.328 | 1.5e-05 | *** |
| cageIDDR_C1 | -0.07742 | 0.08243 | 0.939 | 0.3476 | |
| cageIDDR_C2 | -0.14006 | 0.07306 | 1.917 | 0.0552 | . |
| cageIDDR_DR1 | -0.38949 | 0.08743 | 4.455 | 8.4e-06 | *** |
| cageIDDR_DR2 | -0.16659 | 0.07441 | 2.239 | 0.0252 | * |
| Model-averaged coefficients: females only | | | | | |
| Full average | | | | | |
| cageIDC_C2 | 0.002111 | 0.092872 | 0.023 | 0.9819 | |
| cageIDC_DR1 | -0.107456 | 0.119383 | 0.900 | 0.3681 | |
| cageIDC_DR2 | -0.274199 | 0.144394 | 1.899 | 0.0576 | . |
| cageIDDR_C1 | -0.084467 | 0.101305 | 0.834 | 0.4044 | |
| cageIDDR_C2 | -0.120907 | 0.098277 | 1.230 | 0.2186 | |
| cageIDDR_DR1 | -0.303821 | 0.161252 | 1.884 | 0.0595 | . |
| cageIDDR_DR2 | -0.095970 | 0.095261 | 1.007 | 0.3137 | |
| larv_adultC_DR | -0.088002 | 0.131206 | 0.671 | 0.5024 | |
| larv_adultDR_C | -0.042469 | 0.070713 | 0.601 | 0.5481 | |
| larv_adultDR_DR | -0.065284 | 0.100195 | 0.652 | 0.5147 | |
| **Conditional average female** | | | | | |
| cageIDC_C2 | 0.002463 | 0.100308 | 0.025 | 0.980412 | |
| cageIDC_DR1 | -0.125364 | 0.119927 | 1.045 | 0.295866 | |

| | | | | | |
|---|---|---|---|---|---|
| cageIDC_DR2 | -0.319896 | 0.098519 | 3.247 | 0.001166 | ** |
| cageIDDR_C1 | -0.098544 | 0.102888 | 0.958 | 0.338174 | |
| cageIDDR_C2 | -0.141058 | 0.091792 | 1.537 | 0.124363 | |
| cageIDDR_DR1 | -0.354455 | 0.111302 | 3.185 | 0.001449 | ** |
| cageIDDR_DR2 | -0.111965 | 0.093789 | 1.194 | 0.232557 | |
| larv_adultC_DR | -0.264016 | 0.071947 | 3.670 | 0.000243 | *** |
| larv_adultDR_C | -0.127411 | 0.064645 | 1.971 | 0.048730 | * |
| larv_adultDR_DR | -0.195860 | 0.067407 | 2.906 | 0.003665 | ** |
| Males: N = 1178, $N_{events}$ = 1069 | | | | | |
| Top model | | | | | |
| larv_adultC_DR | -0.31340 | 0.08932 | -3.509 | 0.00045 | *** |
| larv_adultDR_C | -0.11852 | 0.08960 | -1.323 | 0.18588 | |
| larv_adultDR_DR | -0.36169 | 0.09150 | -3.953 | 7.72e-05 | *** |
| Fecundity: age-specific number of eggs. Note: empty terms have been omitted but can be retrieved by running the script (see manuscript for link to GitHub repository) | | | | | |
| Full average | | Adjusted | | | |
| (Intercept) | 129.6244 | 36.9908 | 3.504 | 0.000458 | *** |
| age | -2.4505 | 0.7461 | 3.284 | 0.001022 | ** |
| cageC_C2 | 39.6561 | 33.1744 | 1.195 | 0.231938 | |
| cageC_DR1 | -13.9415 | 51.5091 | 0.271 | 0.786652 | |
| cageC_DR2 | 10.6005 | 51.5091 | 0.206 | 0.836948 | |
| cageDR_C1 | 235.6120 | 52.1254 | 4.520 | 6.20e-06 | *** |
| cageDR_C2 | 319.3923 | 51.2138 | 6.236 | < 2e-16 | *** |
| cageDR_DR1 | 50.6508 | 51.5091 | 0.983 | 0.325442 | |
| cageDR_DR2 | 98.6317 | 51.5091 | 1.915 | 0.055513 | . |
| age:laradC_DR | 0.5133 | 1.0091 | 0.509 | 0.610987 | |
| age:laradDR_C | -4.0953 | 1.0178 | 4.024 | 5.73e-05 | *** |
| age:laradDR_DR | -0.6231 | 1.0091 | 0.617 | 0.536921 | |
| **Conditional average** | | | | | |
| (Intercept) | 129.6244 | 36.9908 | 3.504 | 0.000458 | *** |
| age | -2.4505 | 0.7461 | 3.284 | 0.001022 | ** |

| | | | | | |
|---|---|---|---|---|---|
| cageC_C2 | 39.6561 | 33.1744 | 1.195 | 0.231938 | |
| cageC_DR1 | -13.9415 | 51.5091 | 0.271 | 0.786652 | |
| cageC_DR2 | 10.6005 | 51.5091 | 0.206 | 0.836948 | |
| cageDR_C1 | 235.6120 | 52.1254 | 4.520 | 6.20e-06 | *** |
| cageDR_C2 | 319.3923 | 50.9782 | 6.236 | < 2e-16 | *** |
| cageDR_DR1 | 50.6508 | 51.5091 | 0.983 | 0.325442 | |
| cageDR_DR2 | 98.6317 | 51.5091 | 1.915 | 0.055513 | . |
| age:laradC_DR | 0.5133 | 1.0091 | 0.509 | 0.610987 | |
| age:laradDR_C | -4.0953 | 1.0178 | 4.024 | 5.73e-05 | *** |
| age:laradDR_DR | -0.6231 | 1.0091 | 0.617 | 0.536921 | |
| **Fecundity: per female per 3-hour oviposition period** | | | | | |
| Full average | | | | | |
| (Intercept) | 6.983732 | 1.361403 | 5.130 | 3e-07 | *** |
| age | -0.083712 | 0.026941 | 3.107 | 0.00189 | ** |
| laradC_DR | 0.439817 | 1.318614 | 0.334 | 0.73872 | |
| laradDR_C | 3.442820 | 2.735720 | 1.258 | 0.20822 | |
| laradDR_DR | 1.507011 | 1.661719 | 0.907 | 0.36446 | |
| age:laradC_DR | -0.006939 | 0.028523 | 0.243 | 0.80780 | |
| age:laradDR_C | -0.062376 | 0.052119 | 1.197 | 0.23138 | |
| age:laradDR_DR | -0.019176 | 0.031158 | 0.615 | 0.53826 | |
| **Conditional average** | | | | | |
| (Intercept) | 6.98373 | 1.36140 | 5.130 | 3e-07 | *** |
| age | -0.08371 | 0.02694 | 3.107 | 0.00189 | ** |
| laradC_DR | 0.65628 | 1.56603 | 0.419 | 0.67516 | |
| laradDR_C | 5.13730 | 1.56929 | 3.274 | 0.00106 | ** |
| laradDR_C | 5.13730 | 1.56929 | 3.274 | 0.00106 | ** |
| laradDR_DR | 2.24873 | 1.56603 | 1.436 | 0.15102 | |
| age:laradC_DR | -0.01035 | 0.03433 | 0.302 | 0.76297 | |
| age:laradDR_C | -0.09308 | 0.03458 | 2.692 | 0.00711 | ** |
| age:laradDR_DR | -0.02861 | 0.03433 | 0.833 | 0.40456 | |
